## Supplementary Figures for "Repurposing clemastine to target glioblastoma cell stemness"

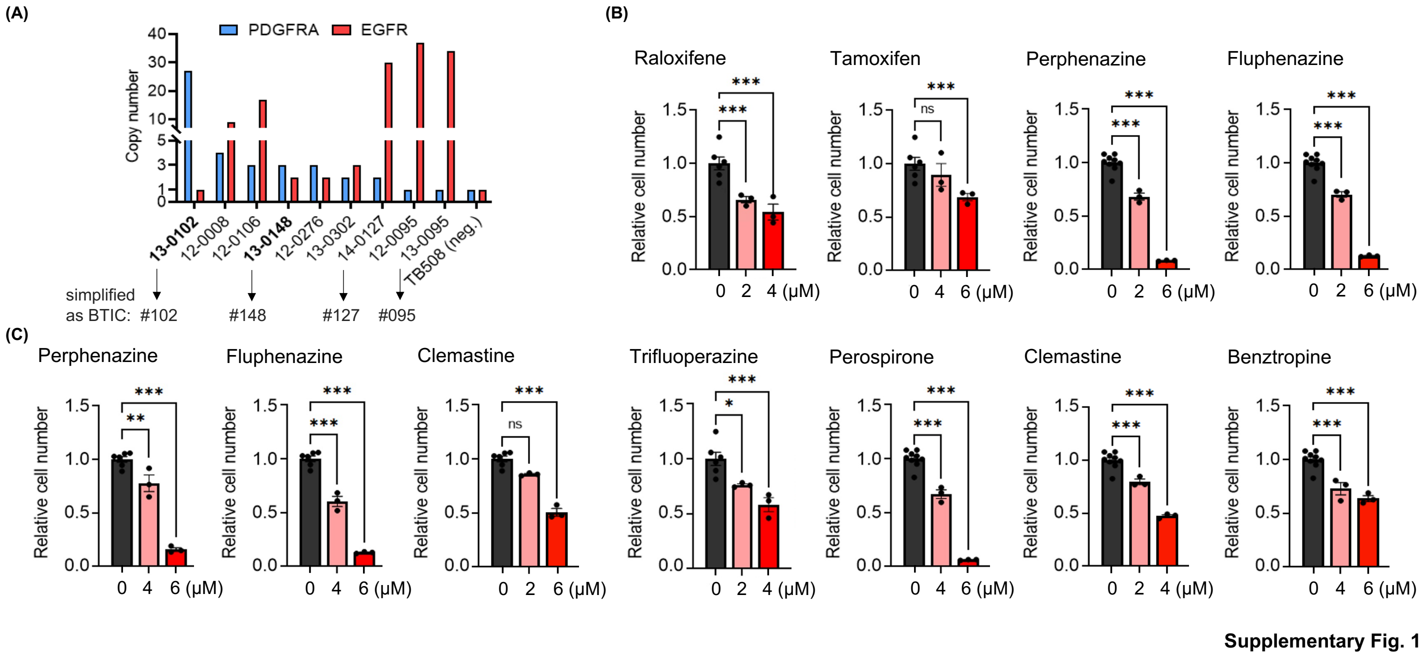


**Supplementary Fig. 1. Patient-derived BTIC cultures bearing *PDGFRA* amplification are susceptible to OPC-differentiating agents. (A)** Normalized copy numbers of *PDGFRA* and *EGFR* genes in 9 patient-derived GBM cultures. Copy numbers of both genes are normalized to the copy numbers of *RNase P*. TB508 serves as a negative control cell line with a single copy of both *PDGFRA* and *EGFR* genes. Simplified IDs for GBM cultures used in this study were noted. **(B-C)** Quantification of relative cell proliferation of **(B)** BTIC#102 and **(C)** BTIC#148 cells treated with indicated OPC-differentiating agents and doses. The y-axis represents normalized phase area confluence at day 7-8 (normalized to day 0 and then normalized to respective vehicle controls). n = 6 for all vehicle groups except for the perospirone, perphenazine, fluphenazine, clemastine, and benztropine-treated BTIC#102 panels, n = 9. n = 3 for all remyelinating agents-treated groups. (B-C) Data are represented as mean ± S.E.M.. Significance was calculated using one-way ANOVA followed by Dunnett's multiple comparisons tests and represented as *p < 0.05, **p < 0.01, ***p < 0.001, n.s.: not significant.

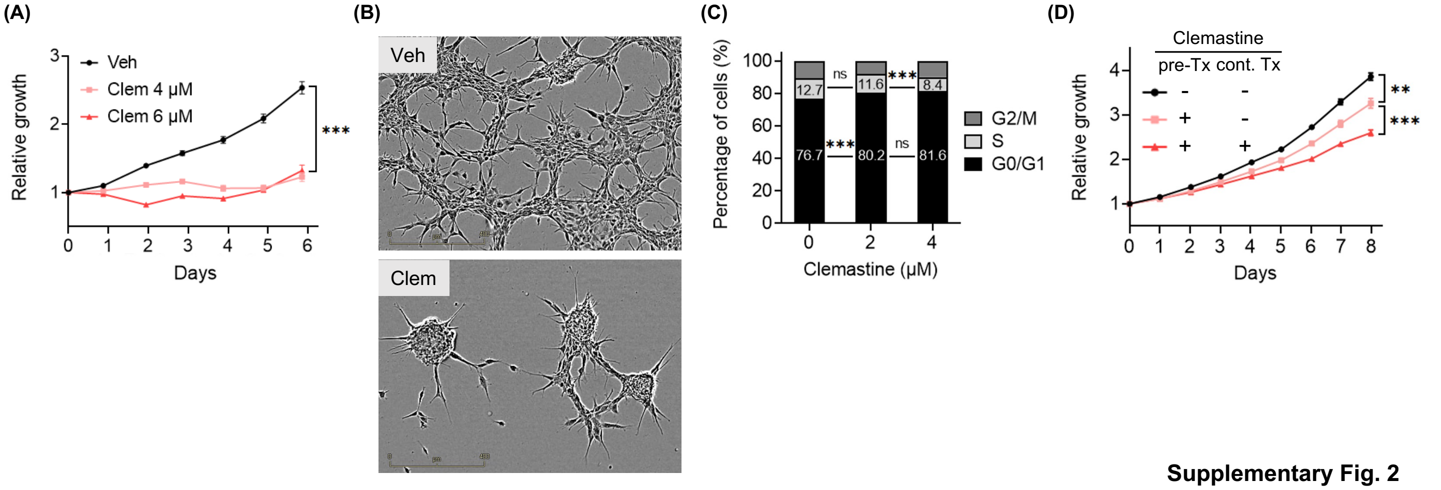


**Supplementary Fig. 2**

**Supplementary Fig. 2. Patient-derived BTIC cultures bearing *PDGFRA* amplification are susceptible to clemastine. (A)** Proliferation of BTIC#148 cells treated with clemastine (Clem) at indicated doses. n = 8 per condition except vehicle, n = 16. **(B)** Representative images (4x, scale bar: 400 μm) of BTIC#148 cells treated with vehicle (Veh) or clemastine (Clem; at 4 μM) for 9 days in laminin-coated plates. **(C)** Quantification of cell cycle analysis of BTIC#148 cells treated with clemastine at indicated doses for 16 days. n = 2 per condition. **(D)** Proliferation of BTIC#148 cells with or without 10-day clemastine (4 μM) pre-treatments (pre-Tx) and/or subsequent clemastine (4 μM) treatments (cont. Tx). Cell proliferation was monitored during subsequent clemastine treatments. n = 18 for pre-$Tx^-$cont.$Tx^-$, n = 9 for pre-$Tx^+$cont.$Tx^-$, and n = 6 for the pre-$Tx^+$cont.$Tx^+$ group. "-": no clemastine in the media; "+": with clemastine in the media. (A, D) Data are represented as mean ± S.E.M.. Significance was calculated using two-way repeated measures ANOVA followed by (A) Dunnett's or (D) Tukey's multiple comparisons tests or (C) two-way ANOVA followed by Tukey's multiple comparisons tests and represented as *$p < 0.05$, **$p < 0.01$, ***$p < 0.001$, n.s.: not significant.

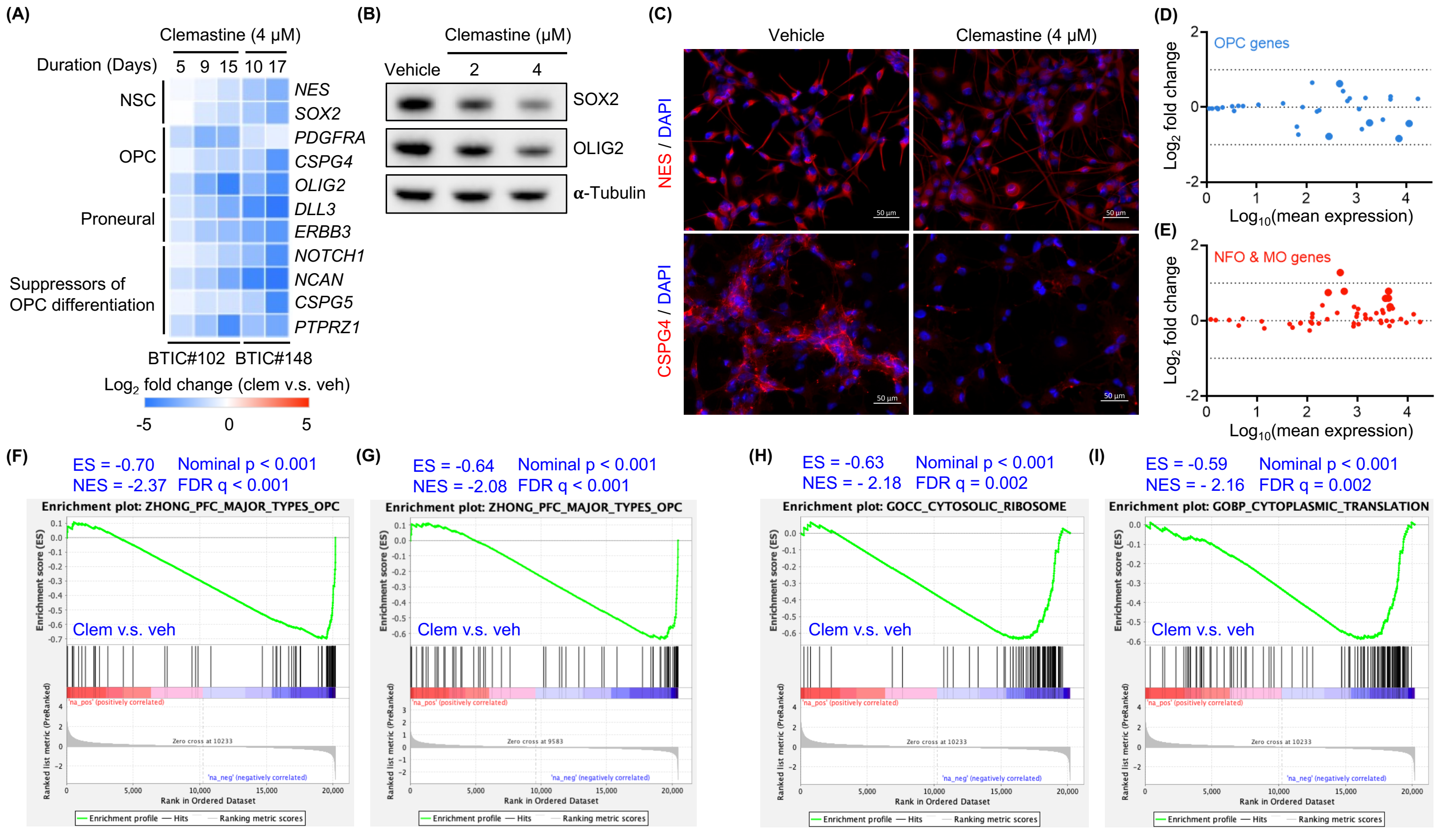


**Supplementary Fig. 3**

**Supplementary Fig. 3. Clemastine treatment attenuates the stemness and progenitor cell features of *PDGFRA*+ BTIC cells. (A)** The heatmap of mRNA expression fold change (clemastine v.s. vehicle) of genes associated with NSCs, OPCs, proneural subtype GBM, and suppressors of OPC differentiation in BTIC#102 and BTIC#148 cells treated with clemastine (4 μM) for the indicated duration of time as assessed by quantitative RT-PCR. n = 3 replicates per condition. **(B-C)** Protein levels of NSC and OPC markers in BTIC#148 cells treated with clemastine at indicated doses for **(B)** 10 days or **(C)** 16 days as assessed by **(B)** immunoblot assays (representative results from two independent experiments are shown) or **(C)** IF staining (representative images from two independent experiments are shown; 20x, scale bar: 50 μm). **(D-E)** MA plots summarizing differential mRNA expression of top 40 **(D)** OPC-specific genes or **(E)** oligodendrocytes-specific genes between clemastine versus vehicle-treated (17-day treatment) BTIC#148 cells. Larger symbols indicate genes with adjusted p-values < 0.05. **(F-I)** GSEA plots comparing the transcriptomic profiles between clemastine (Clem) versus vehicle (Veh)-treated **(F, H-I)** BTIC#102 (15-day treatment) or **(G)** BTIC#148 (17-day treatment) cells with the following gene sets: **(F-G)** OPC, (**H**) cytosolic ribosome, and **(I)** cytoplasmic translation gene sets.

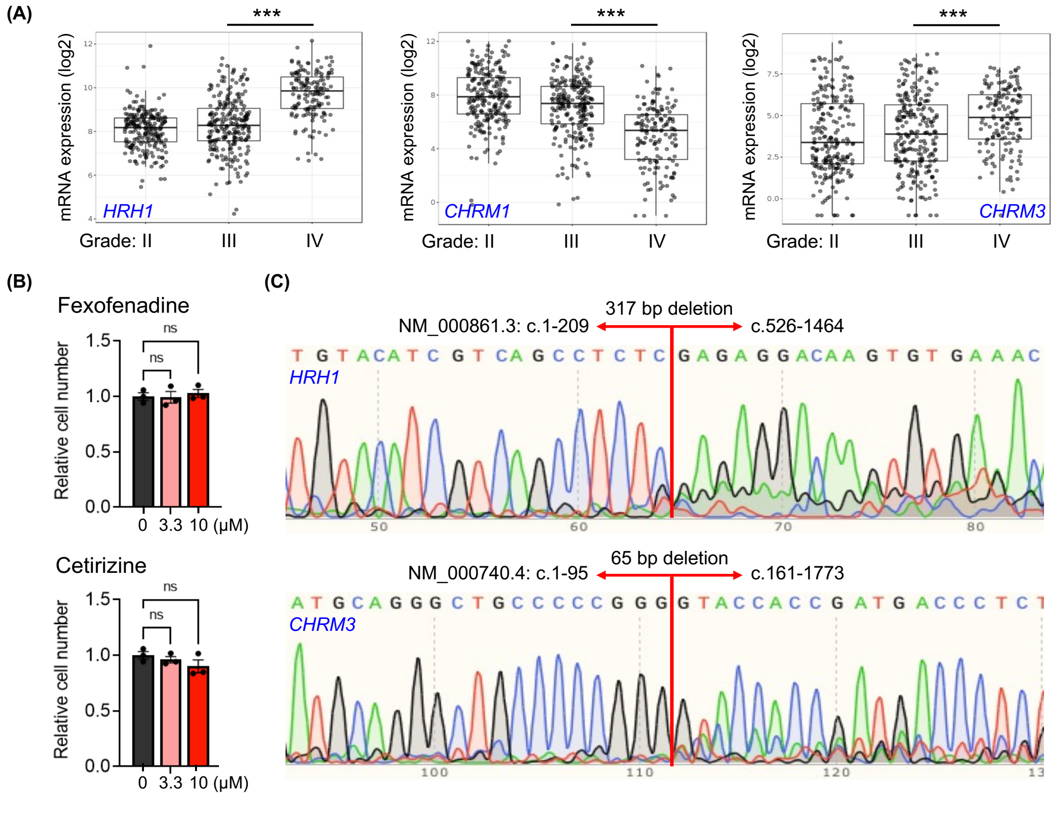


Supplementary Fig. 4

**Supplementary Fig. 4. Assessing the roles of potential pharmacological targets of clemastine in BTICs. (A)** Boxplots of mRNA expression levels of *HRH1*, *CHRM1*, and *CHRM3* in GBMs and low-grade gliomas (data retrieved from TCGA_GBMLGG dataset). Plots were generated using the GlioVis portal. **(B)** Quantification of relative cell proliferation of BTIC#102 cells treated with indicated H1R antagonists and doses. The y-axis represents normalized phase area confluence at day 7 (normalized to day 0 and then normalized to respective vehicle controls). n = 3 per condition. **(C)** Sanger sequencing results of the amplicon targeting the coding sequence of *HRH1* gene in *HRH1*-knockout BTIC#102 cells (upper panel) and *CHRM3* gene in *CHRM3*-knockout BTIC#102 cells (lower panel). The results indicate a 317 bp deletion in the coding region of the *HRH1* gene (c.210_525del) and a 65 bp deletion in the coding region of the *CHRM3* gene (c.96_160del). **(D)** Proliferation of non-targeting sgControl (sgCtl) and *HRH1*-knockout (*HRH1*-KO) BTIC#102 cells (upper panel) or *CHRM3*-knockout (*CHRM3*-KO) BTIC#102 cells (lower panel) treated with vehicle (Veh) or clemastine (Clem; at 4 µM). n = 3 per condition. (B, D) Data are represented as mean ± S.E.M.. Significance was calculated using (A) Tukey's HSD tests by the GlioVis portal, (B) one-way ANOVA followed by Dunnett's multiple comparisons tests, or (D) two-way repeated measures ANOVA followed by Tukey's multiple comparisons tests, and represented as *p < 0.05, **p < 0.01, ***p < 0.001, n.s.: not significant.

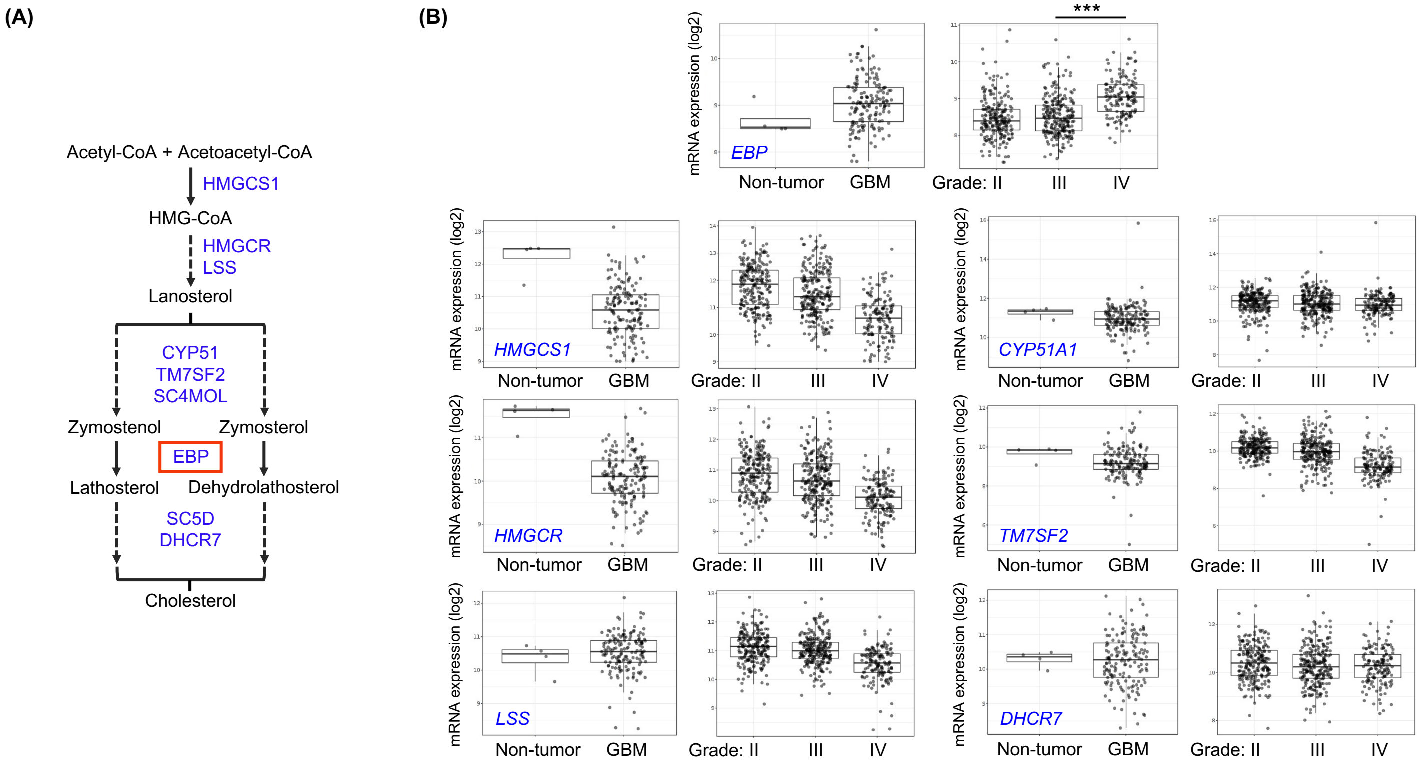


**Supplementary Fig. 5**

**Supplementary Fig. 5. *EBP* displays distinct expression patterns in gliomas in comparison to other genes in the cholesterol biosynthesis pathway. (A)** Schematic representation of sterol metabolites (in black) and enzymes (in blue) involved in the human cholesterol biosynthesis pathway. **(B)** Boxplots of mRNA expression levels of genes in the cholesterol biosynthesis pathway (data for *SC4MOL* and *SC5D* were not available) in GBMs and non-tumors (left panel, data retrieved from TCGA_GBM dataset and RNA-seq platform) or low-grade gliomas (right panel, data retrieved from TCGA_GBMLGG dataset). Plots were generated using the GlioVis portal. Significance was calculated using Tukey's HSD tests by the GlioVis portal and represented as *p < 0.05, **p < 0.01, ***p < 0.001, n.s.: not significant.

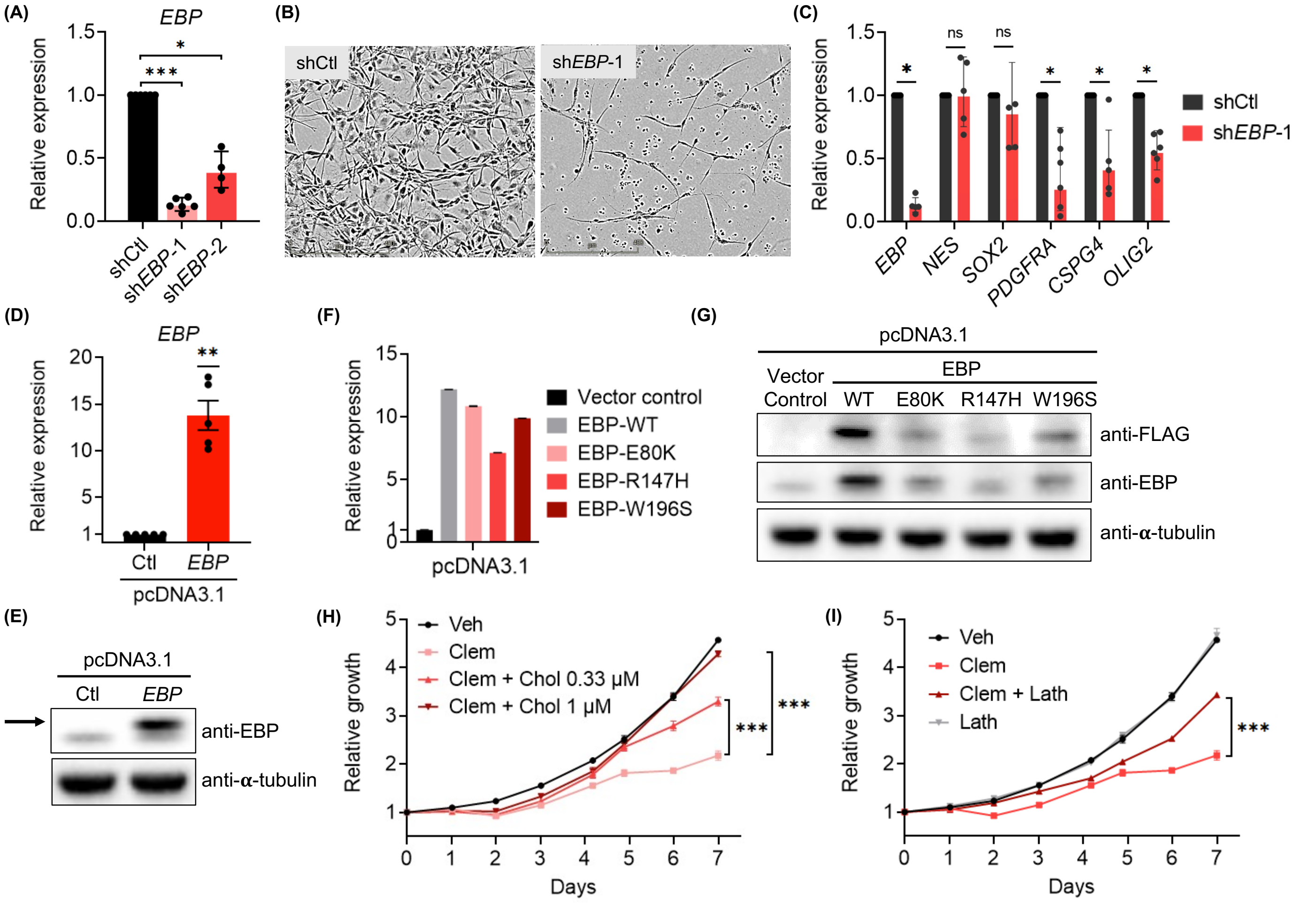


**Supplementary Fig. 6**

**Supplementary Fig. 6. EBP is essential for BTIC propagation. (A)** mRNA expression levels of *EBP* in shCtl, sh*EBP*-1, and sh*EBP*-2 BTIC#102 cells as assessed by quantitative RT-PCR. n = 6 per condition except sh*EBP*-2, n = 4. **(B)** Representative images (4x, scale bar: 400 μm) of shCtl and sh*EBP*-1 BTIC#148 cells 10 days after lentiviral transduction. **(C)** mRNA expression levels of genes associated with NSCs and OPCs in shCtl and sh*EBP*-1 BTIC#102 cells as assessed by quantitative RT-PCR. n = 5 per gene except *PDGFRA* and *OLIG2*, n = 6. **(D)** mRNA expression levels of *EBP* in vector control (pcDNA3.1-Ctl) and *EBP*-overexpressed (pcDNA3.1-*EBP*) BTIC#102 cells. n = 5 per condition. **(E)** Protein levels of EBP in vector control and *EBP*-overexpressed BTIC#102 cells as assessed by immunoblot assays (representative results from two independent experiments are shown). **(F)** mRNA expression levels of *EBP* (wild-type or mutant) in BTIC#102 cells with vector control, overexpression of wild-type EBP, or overexpression of each of the three mutants as assessed by quantitative PCR. n = 3 replicates per group. **(G)** Protein levels of FLAG-EBP (wild-type or mutant) in BTIC#102 cells with vector control, overexpression of wild-type EBP, or overexpression of each of the three mutants as assessed by immunoblot assays (representative results from two independent experiments are shown) using anti-FLAG and anti-EBP antibodies. **(H)** Proliferation of BTIC#148 cells treated with clemastine (Clem; at 6 μM) with or without water-soluble cholesterol (Chol) at indicated doses. n = 6 per condition. **(I)** Proliferation of BTIC#148 cells treated with clemastine (Clem; at 6 μM) and/or lathosterol (Lath; at 3.13 μM). n = 6 per condition. (A, C-D) Data are represented as geometric mean ± geometric S.D. of fold change relative to control groups, (F) geometric mean ± S.E.M., or (H-I) mean ± S.E.M.. Significance was calculated using (H-I) two-way repeated measures ANOVA followed by Tukey's multiple comparisons tests and represented as *$p < 0.05$, **$p < 0.01$, ***$p < 0.001$, n.s.: not significant.

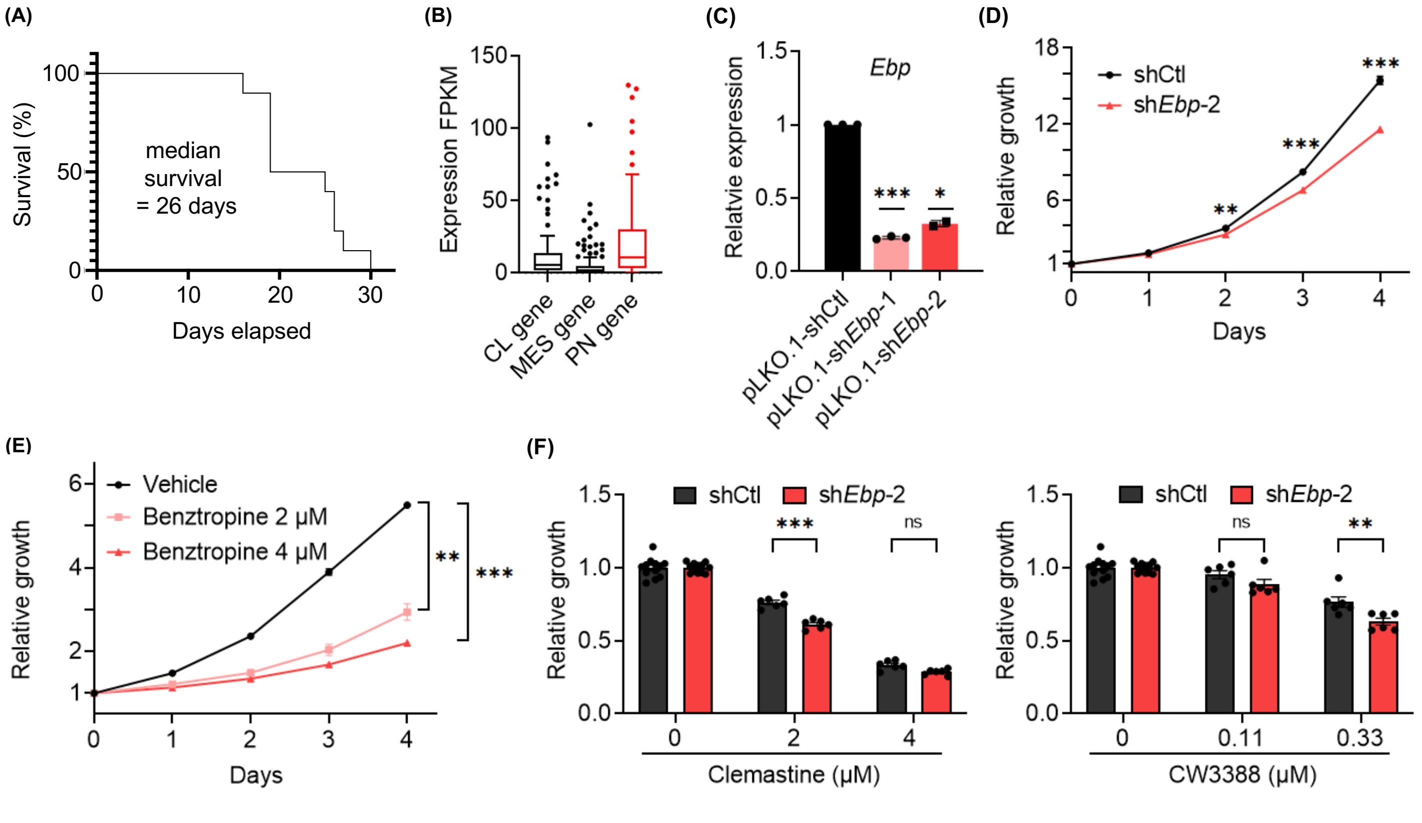


**Supplementary Fig. 7**

**Supplementary Fig. 7. Loss of Ebp impairs the growth of a mouse glioma cell line (C266) derived from mouse p53-null NSCs with PDGFB overexpression. (A)** The Kaplan Meier analyses of mice orthotopically implanted with mouse p53-null NSCs transduced with PDGFB-overexpressing retrovirus. n = 10 mice. **(B)** Boxplots of mRNA expression levels (FPKM) of markers genes for classical (CL, n = 127), mesenchymal (MES, n = 174), or proneural (PN, n = 133) in mouse glioma tumors as assessed by mRNA-seq (only genes detected in the tumor samples and matched between mouse and human were used for the analysis. Note the intra-tumoral heterogeneity and the relative dominance of the PN subtype). An independent analysis using the human gene training set described in Ref. 84 yielded consistent subtyping results. **(C)** Quantitative RT-PCR analyses confirmed the reduced *Ebp* transcript levels induced by gene knockdown. n = 3 per condition except sh*Ebp*-2, n = 2. **(D)** Proliferation of shCtl and sh*Ebp*-2 C266 cells. P-values indicate significance between shCtl and sh*Ebp*-2 cells at indicated timepoints. n = 12 per condition. **(E)** Proliferation of C266 cells treated with benztropine at indicated doses. n = 6 per condition except vehicle, n = 12. **(F)** Quantification of relative cell proliferation of shCtl and sh*Ebp-2* C266 cells treated with clemastine (left panel) or CW3388 (right panel) at indicated doses. The y-axis represents normalized phase area confluence at day 4 (normalized to day 0 and then normalized to respective vehicle controls). n = 6 per condition except for the vehicle groups of the clemastine-treated panel, n = 9. Data are represented as (C) geometric mean ± geometric S.D. of fold change relative to the shCtl group or (D-F) mean ± S.E.M.. Significance was calculated using two-way repeated measures ANOVA followed by (D) Sidak's or (E) Dunnett's multiple comparisons tests, or (F) two-way ANOVA followed by Sidak's multiple comparisons tests, and represented as *$p < 0.05$, **$p < 0.01$, ***$p < 0.001$, n.s.: not significant.

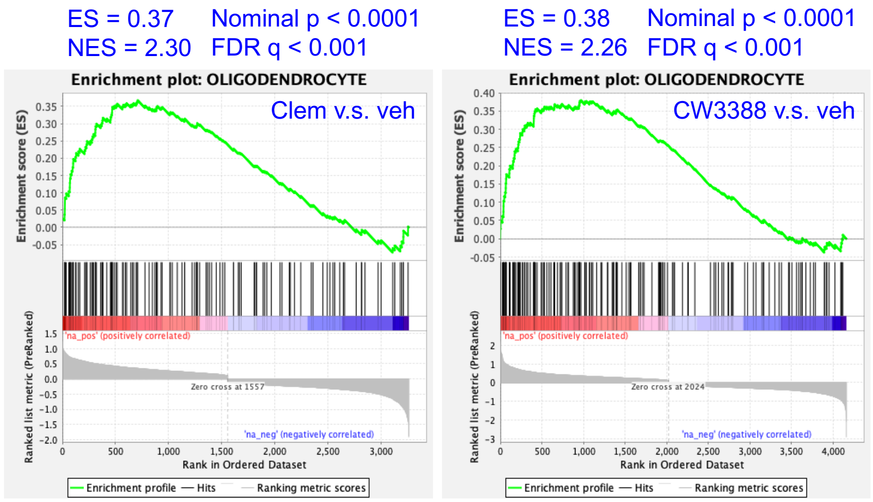


**Supplementary Fig. 8**

**Supplementary Fig. 8. Clemastine treatment broadly alters multiple signaling pathways in a mouse glioma cell line (C266).** GSEA plots of genes differentially expressed between (left panel) clemastine (Clem) or (right panel) CW3388 versus vehicle (Veh)-treated C266 cells (12-day treatment) with the "OLIGODENDROCYTE" gene set (includes top 40 oligodendrocyte-specific genes).
