## Supplementary Materials for "Repurposing clemastine to target glioblastoma cell stemness"

| RT-qPCR primer sequences: | | |
|---|---|---|
| Gene name | Forward (F) or Reverse (R) | Sequence (5' to 3') |
| *ACTB* | F | GATTCCTATGTGGGCGACGA |
| | R | AGGTCTCAAACATGATCTGGGT |
| *B2M* | F | TGCCGTGTGAACCATGTG |
| | R | ACCTCCATGATGCTGCTTACA |
| *NES* | F | CTGGAGCAGGAGAAACAGGG |
| | R | GAGGGAAGTCTTGGAGCCAC |
| *SOX2* | F | AACGGCAGCTACAGCATGAT |
| | R | GACTTGACCACCGAACCCAT |
| *PDGFRA* | F | GACGGTCTTGGAAGTGAGCA |
| | R | TAAAGCCCTGTCTGCTGTCG |
| *CSPG4* | F | GGGCCGTTCTCTATAGCCAC |
| | R | GACACCATCACCAGGTAGCC |
| *OLIG2* | F | AGTGGCTTCAAGTCATCCTCG |
| | R | TGTTGATCTTGAGACGCAGCT |
| *DLL3* | F | TAGCGCTCATTTTCCTCCCC |
| | R | TGGATCTGCAGCTCGAAGAC |
| *PTPRZ1* | F | GCAGTTGGATGGAGAGGACC |
| | R | TCGACAATCAGTTGGTCGCT |
| *CSPG5* | F | GAGCTTCTAGTGCCCACTGG |
| | R | CATCCCCTTGTGCCAGATGT |
| *NOTCH1* | F | CAGATCCTGATCCGGAACCG |
| | R | CCAGAAACAGGGGTGTCTCC |
| *NCAN* | F | GTGTCACTGCCTTCCTACCC |
| | R | TGCAATGATGGCTGAGCTGA |
| *MOG* | F | AGGGAAAGGTGACTCTCAGGA |
| | R | GGAAGATGAGGCCAACAGTGA |
| *ERBB3* | F | TGAATGGCCTGAGTGTGACC |
| | R | ATCGTAGACCTGGGTCCCTC |
| *EBP* | F | TACTGGCTGGCCTCTTCTCT |
| | R | CCCAGGATGTATCGGCTGTC |
| *Actb* | F | GGTGGGAATGGGTCAGAAGG |
| | R | GTACATGGCTGGGGTGTTGA |
| *B2m* | F | TTTCAGTGGCTGCTACTCGG |
| | R | TGTTCGGCTTCCCATTCTCC |

| | | |
|---|---|---|
| *Ebp* | F | ATCGAGGGCTGGTTCTCTCT |
| | R | GCCCATAGACACCACAAGCT |

| **shRNA sequences:** | | |
|---|---|---|
| Name | Clone ID | Sequence (5’ to 3’) |
| *EBP* #1 (human) | TRCN0000049393 | CCGG-GCACCTAAGACTGGACAACTT-CTCGAG-AAGTTGTCCAGTCTTAGGTGC-TTTTTG |
| *EBP* #2 (human) | TRCN0000049394 | CCGG-CTCCGCTTCATTCTACAGCTT-CTCGAG-AAGCTGTAGAATGAAGCGGAG-TTTTTG |
| *Ebp* #1 (mouse) | TRCN0000111870 | CCGG-CCAGAAGACTCAAATCTTCTT-CTCGAG-AAGAAGATTTGAGTCTTCTGG-TTTTTG |
| *Ebp* #2 (mouse) | TRCN0000111872 | CCGG-CCAAGGGAGATAGCCGATATA-CTCGAG-TATATCGGCTATCTCCCTTGG-TTTTTG |
| Non-targeting control | SHC002 | CCGG-CAACAAGATGAAGAGCACCAA-CTCGAG-TTGGTGCTCTTCATCTTGTTG-TTTTTG |

| **sgRNA sequences:** | | |
|---|---|---|
| Name | Forward (F) or Reverse (R) | Sequence (5’ to 3’) |
| sg-*HRH1*-exon2-F1 | F | CACCGTACATCGTCAGCCTCTCGG |
| sg-*HRH1*-exon2-R1 | R | AAACCCGAGAGGCTGACGATGTAC |
| sg-*HRH1*-exon2-F2 | F | CACCGCAGACCTCGGTGCGCCGAG |
| sg-*HRH1*-exon2-R2 | R | AAACCTCGGCGCACCGAGGTCTGC |
| sg-*CHRM3*-exon5-F1 | F | CACCGGGTCATCGGTGGTACCGTC |
| sg-*CHRM3*-exon5-R1 | R | AAACGACGGTACCACCGATGACCC |
| sg-*CHRM3*-exon5-F2 | F | CACCGAAATGAGTGACGGTTCCCG |
| sg-*CHRM3*-exon5-R2 | R | AAACCGGGAACCGTCACTCATTTC |
| sgRNA-control-F | F | CACCGTAGGCGCGCCGCTCTCTAC |
| sgRNA-control-R | R | AAACGTAGAGAGCGGCGCGCCTAC |

| **PCR primer sequences:** | | |
|---|---|---|
| Name | Forward (F) or Reverse (R) | Sequence (5’ to 3’) |
| PCR-*HRH1*-F1 | F | GGTCACAGTAGGGCTCAACC |
| PCR-*HRH1*-R1 | R | CACCACCAGCATCTTTTGGC |
| PCR-M13F-*CHRM3*-F1 | F | TGTAAAACGACGGCCAGTCCCCCAGACTATGTCAGAGAG |

| | | |
|---|---|---|
| PCR-M13F-*CHRM3*-R1 | R | CAGGAAACAGCTATGACCCTAAGGCCCATCGATT CATGA |
| pLentiCRISPR-R1 | N/A | CCCGTTGCGAAAAAGAACGT |

| ***EBP* mutagenesis primer sequences:** | | |
|---|---|---|
| EBP mutation | Forward (F) or Reverse (R) | Sequence (5' to 3') |
| E80K | F | CATTCACCTGGTGATCAAGGGCTGGTTCGTTCT |
| | R | AGAACGAACCAGCCCTTGATCACCAGGTGAATG |
| R147H | F | GCCAGCATCCCCTCCACTTCATTCTACAGCT |
| | R | AGCTGTAGAATGAAGTGGAGGGGATGCTGGC |
| W196S | F | GTCTTCATGAATGCCCTGTCGCTGGTGCTG |
| | R | CAGCACCAGCGACAGGGCATTCATGAAGAC |

| **Plasmids:** | | |
|---|---|---|
| Name | Provider | Cat. Number |
| pcDNA3.1 V5-His A | GenScript | N/A |
| pcDNA3.1$^+$ C-(K)-DYK-hEBP | GenScript | OHu18817 |
| pcDNA3.1$^+$ C-(K)-DYK-hEBP-E80K | This lab | N/A |
| pcDNA3.1$^+$ C-(K)-DYK-hEBP-R147H | This lab | N/A |
| pcDNA3.1$^+$ C-(K)-DYK-hEBP-W196S | This lab | N/A |
| EGFP-hGal3 | Addgene | 73080 |
| PM-GFP | Addgene | 21213 |
| LentiCRISPRv2E | Addgene | 78852 |

| **Chemical compounds:** | | |
|---|---|---|
| Name | Vendor | Cat. Number |
| Clemastine (fumarate) | Cayman | 14637 |
| Lathosterol | Cayman | 9003102 |
| Cholesterol-water soluble | Sigma-Aldrich | C4951 |
| Tamoxifen | Sigma-Aldrich | T5648 |
| Puromycin dihydrochloride | Sigma-Aldrich | P8833 |
| Dimethyl sulfoxide | Sigma-Aldrich | D2650-100ML |
| Ascorbic acid | Lonza | CC-4398 |
| Geneticin (G418 Sulfate) | ThermoFisher | 10131035 |
| CW3388 | Sigma-Aldrich | SML2602 |
| Polybrene | Sigma-Aldrich | TR-1003 |

| | | |
|---|---|---|
| TASIN-1 | Cayman | 21155 |
| Cetirizine | Cayman | 19686 |
| Fexofenadine | Cayman | 18191 |
| D-luciferin, sodium salt | Goldbio | LUCNA-1 |
| 2-Mercaptoethanol | Sigma-Aldrich | M7522-100ML |
| Tween 20 | BIO-RAD | 1706531 |
| Methanol | VWR | BDH1135-4LP |
| Ethanol absolute | VWR | 89125-172 |
| Isoflurane | Patterson | 07-893-8441 |
| DAPI | MilliporeSigma | D9542-10MG |
| Fluphenazine (hydrochloride) | Cayman | 23555 |
| Perphenazine | Cayman | 20735 |
| Trifluoperazine (hydrochloride) | Cayman | 15068 |
| Raloxifene (hydrochloride) | Cayman | 10011620 |
| Benztropine (mesylate) | Cayman | 16214 |
| Saponin | MilliporeSigma | 84510-100G |

| **Reagents and kits:** | | |
|---|---|---|
| Name | Vendor | Cat. Number |
| Human PDGF-AA | Shenandoah | 100-16 |
| Human PDGF-BB | Shenandoah | 100-18 |
| Laminin | MilliporeSigma | L2020 |
| Triton X-100 solution | Fluka | 93443 |
| Formalin 10%, neutral buffered | VWR | 89370-094 |
| Pierce™ 16% Formaldehyde (w/v), Methanol-free | ThermoFisher | 28908 |
| Ponceau S solution | Sigma-Aldrich | P7170 |
| Nitrocellulose Membrane, 0.45um; Roll 30 cm x 3.5 m | Bio-Rad | 1620115 |

| | | |
|---|---|---|
| Trans-Blot Turbo RTA Mini 0.2 µm Nitrocellulose Transfer Kit | Bio-Rad | 1704270 |
| NuPAGE 4-12% Bis-Tris Protein Gels, 1.0mm, 12 well | ThermoFisher | NP0322BOX |
| NuPAGE 4-12% Bis-Tris Protein Gels, 1.0mm, 10 well | ThermoFisher | NP0321BOX |
| NuPAGE 4-12% Bis-Tris Protein Gels, 1.5mm, 15 well | ThermoFisher | NP0336BOX |
| NuPAGE 4-12% Bis-Tris Protein Gels, 1.5mm, 10 well | ThermoFisher | NP0335BOX |
| Mini Gel Tank | ThermoFisher | A25977 |
| RNA to cDNA EcoDry™ Premix (Double Primed) | Takara (Clontech) | 639548 |
| KAPA SYBR FAST qPCR Master Mix (2X) | Roche | KK4602 |
| SuperSignal™ West Pico PLUS Chemiluminescent Substrate | ThermoFisher | 34580 |
| Pierce™ BCA Protein Assay Kit | ThermoFisher | 23225 |
| Accutase solution | MilliporeSigma | A6964-500ML |
| Corning 96-well Clear Flat Bottom Polystyrene TC-treated Microplate | Corning | 3595 |
| Quick-RNA miniprep kit | Genesee | 11-328 (Zymo R1055) |
| UltraPure DNase/RNase-Free Distilled Water | ThermoFisher | 10977015 |
| Dulbecco's Phosphate Buffered Saline | MilliporeSigma | D8537-500ML |
| PBS, pH 7.4 | ThermoFisher | 10010023 |
| DPBS, no calcium, no magnesium | ThermoFisher | 14190144 |
| 4X Laemmli Sample Buffer | BIO-RAD | 1610747 |
| Trypan Blue Solution (0.4%) | Sigma-Aldrich | T8154 |
| Opti-MEM™ I Reduced Serum Medium | ThermoFisher | 31985070 |
| McCoy's 5A (Modified) Medium | ThermoFisher | 16600082 |
| Trypsin (2.5%), no phenol red | ThermoFisher | 15090046 |
| Lipofectamine 3000 Transfection Reagent | ThermoFisher | L3000008 |
| Bovine Serum Albumin | Sigma-Aldrich | A9647-100G |
| Restore Western Blot Stripping Buffer | ThermoFisher | 21059 |
| NuPAGE MOPS SDS Running Buffer (20X) | ThermoFisher | NP0001 |

| | | |
|---|---|---|
| NuPAGE Antioxidant | ThermoFisher | NP0005 |
| Corning 1L 10X Tris Buffered Saline | Corning | 46-012-CM |
| 10x Tris Buffered Saline (TBS) | BIO-RAD | 1706435 |
| ProLong Gold Antifade Mountant | ThermoFisher | P36934 |
| SlowFade Diamond Antifade Mountant | ThermoFisher | S36972 |
| FxCycle™ PI/RNase Staining Solution | ThermoFisher | F10797 |
| S.O.C. Medium | ThermoFisher | 15544034 |
| TempPlate full-skirted 96-well PCR plate | USA Scientific | 1402-9800 |
| Precision Plus Protein Dual Color Standards | BIO-RAD | 1610374 |
| 50x TAE (Tris/Acetic Acid/EDTA) Buffer, 1 L | BIO-RAD | 1610743 |
| Non-fat dry milk | Genesee | 20-241 |
| Goat serum | MilliporeSigma | G9023-10ML |
| QuikChange Lightning Multi Site-Directed Mutagenesis Kit | Agilent | 210519 |
| ZymoPURE Plasmid Miniprep Kit | Genesee | 11-553 |
| Neon Transfection System 10 µL Kit | ThermoFisher | MPK1025 |

| **Equipment:** | | |
|---|---|---|
| Name | Vendor | Cat. Number |
| Bioruptor Standard | Diagenode | UCD-200 |
| CFX96 Real-Time PCR Detection System | BIO-RAD | N/A |
| Nunc Lab-Tek II Chamber Slide System - 2-well Chamber Slide w/ removable wells | ThermoFisher | 154461 |
| PowerPac Basic Power Supply | BIO-RAD | 1645050 |
| Trans-Blot Turbo Transfer System | BIO-RAD | 1704150 |
| ChemiDoc MP System | BIO-RAD | 1708280 |

| **Antibody:** | | | | | |
|---|---|---|---|---|---|
| Name | Vendor | Cat. Number | RRID | Dilution for blotting | Dilution for IF |
| Nestin | MilliporeSigma | ABD69 | AB_2744681 | | **1:500** |
| SOX2 | Cell Signaling | 3579S | AB_2195767 | **1:1000** | **1:400** |
| PDGF receptor **α** | Cell Signaling | 3174S | AB_2162345 | **1:1000** | |
| PDGF receptor **α** | Cell Signaling | 5241S | AB_10692773 | | **1:400** |

| | | | | | |
|---|---|---|---|---|---|
| CSPG4 | Cell Signaling | 43916S | N/A | | **1:200** |
| OLIG2 | Novus | NBP1-28667 | AB_1914109 | **1:3000** | **1:1000** |
| Galactocerebroside | MilliporeSigma | MAB342 | AB_94857 | | **1:200** |
| EBP | Santa Cruz | sc-374267 | AB_11012162 | **1:500** | |
| $\alpha$-tubulin | Cell Signaling | 3873S | AB_1904178 | **1:1000** | |
| Anti-rabbit IgG, HRP-linked Antibody | Cell Signaling | 7074S | AB_2099233 | **1:1000** | |
| Anti-mouse IgG, HRP-linked Antibody | Cell Signaling | 7076S | AB_330924 | **1:1000** | |
| Goat anti-Rabbit IgG (H+L) - Alexa Fluor 594 | ThermoFisher | A-11037 | AB_2534095 | | **1:500** |
| Goat anti-Rabbit IgG (H+L) - Alexa Fluor 647 | ThermoFisher | A-21245 | AB_2535813 | | **1:500** |
| Goat anti-Rabbit IgG (H+L) - Alexa Fluor Plus 488 | ThermoFisher | A32731 | AB_2633280 | | **1:500** |
| Goat anti-Mouse IgG (H+L) - Alexa Fluor 488 | ThermoFisher | A-11029 | AB_2534088 | | **1:500** |
| Goat anti-Mouse IgG (H+L) - Alexa Fluor Plus 488 | ThermoFisher | A32723 | AB_2633275 | | **1:500** |
| Goat anti-Mouse IgG (H+L) - Alexa Fluor Plus 647 | ThermoFisher | A32728 | AB_2633277 | | **1:500** |
| Normal Rabbit IgG | MilliporeSigma | 12-370 | AB_145841 | | **1:800-1:1000** |
| Normal Mouse IgG | MilliporeSigma | 12-371 | AB_145840 | | **1:200-1:1000** |
| Rabbit IgG Isotype Control | ThermoFisher | 31235 | AB_243593 | | **1:1200-1:20000** |
| Mouse IgG Isotype Control | ThermoFisher | 31903 | AB_10959891 | | **1:2500** |
